# High-throughput screening of a *Debaryomyces hansenii* library for potential candidates with improved stress tolerance and wider carbon utilisation capabilities

**DOI:** 10.1101/2022.03.24.485636

**Authors:** Anne Sofie B. Dyerberg, Clara Navarrete, José L. Martínez

## Abstract

Industrial biotechnology comprises the manufacturing of bulk chemicals and high-value end-products from renewable feedstocks, thus it presents a valuable aspect in the present transition from traditional-resource demanding manufacturing to sustainable solutions. The non-conventional yeast *Debaryomyces hansenii* encompasses halotolerant characteristics that ensures its use in industrial applications, and hence, its industrial importance. For this purpose, a comprehensive and holistic understanding of its behaviour and response to abiotic stresses is essential. Through high-throughput screening methods, using advanced robotics and automation devices, the present study enlightens intraspecies behavioural characteristics of novel *D. hansenii* strains in response to sodium, as well as their ability to tolerate abiotic stress in semi-controlled micro-fermentations and spot-test studies. A significantly improved performance under those abiotic stresses was observed under the presence of 1M NaCl. Moreover, a positive and summative effect on growth was also found in pH 4 and high salt content. Our results align with previous findings suggesting the halophilic (and not just halotolerant) behaviour of *D. hansenii*, which is now extensive to all the *D. hansenii* strains included in this study. Strain-specific differential responses to the presence of sodium were also observed, with some strains exerting a more notable induction by the presence of salt than the standard strain (CBS767). Furthermore, our study provides indications of the use of *D. hansenii* in industrial bioprocesses based on lignocellulosic biomass and non-lignocellulosic feedstocks.

## Introduction

Industrial biotechnology comprises the ability to manufacture bulk chemicals and high-value products from renewable feedstocks. Rapid increase in innovative industrial biotech tools have reduced cost of production, thus creating competitive advantages for the production of biomass-derived products and making them commercially successful (Li et al. 2014). However, several hurdles still need to be addressed to fully realise and obtain the comprehensive potential of industrial bioprocessing (Erickson, Nelson, and Winters 2012). Enabling high yield and reduced manufacturing maintenance, including reduced sterilisation processes, remain a current obstacle in manufacturing biotechnological products and it reflects the often-costly processes. Deficiency of cost-effective solutions decelerate the diversion towards a bio-based economy, hence continued research in optimum hosts and processes is needed to maximise commercial success and competitive methods (Li et al. 2014). Saline environments permit reduced sterilisation costs due to their natural barrier toward microbial contaminants, for which the high salinity is toxic. Therefore, using saline environments in industrial bioprocesses offer the potential of minimising operational costs (Navarrete et al. 2020b).

The hemiascomycetous non-conventional yeast *Debaryomyces hansenii* encompasses physiological properties that have generated an increasing interest in the yeast (Prista et al. 2016). Its unique and prominent halophilic/halotolerant characteristics ensure its industrial applications, and hence, its industrial importance. It is described to be able to tolerate levels up to 4.11 M of sodium (Chao, Yen, and Ku 2009). However, growth inhibition has been observed at concentrations exceeding 2M NaCl (Navarrete et al. 2020a; Capusoni et al. 2019). Additionally, when *D. hansenii* is grown in 1M NaCl, protective and non-detrimental effects have been observed. Presence of stress factors, including extreme pH conditions, oxidative stress or different temperatures, are proved to have less detrimental effect when in co-existence with NaCl (Prista et al. 2016). At high temperatures with severe growth inhibitions, presence of salt has also appeared to stimulate growth of the yeast. Additionally, stress by extreme pH has shown to be relieved by the presence of 0.25M NaCl (Almagro et al. 2000). Similar results were obtained by other authors, observing a survival strategy of *D. hansenii* in high salinity environments (Butinar, Strmole, and Gunde-Cimerman 2011; Navarrete et al. 2020a).

Furthermore, *D. hansenii* can utilize a broad spectrum of carbon substrates, generally catabolised through respiratory metabolism (Nobre, Lucas, and Leão 1999; Palma et al. 2007). Industrial substrates often contain a variety of lignocellulosic biomass and non-lignocellulosic feedstocks, which are composed of mixtures of hexoses and pentoses (Navarrete et al. 2020b). Hence, *D. hansenii’*s ability to assimilate different carbon sources encourages further research about its potential applications in industrial bioprocesses (Prista et al. 2016).

In 2016, Prista *et al*. reviewed the main characteristics of *D. hansenii* that makes the use of this yeast interesting for food and biotechnology applications. Moreover, there is a demand for studies of wild *D. hansenii*-strains isolated from natural habitats, to increase the understanding of this yeast, while enabling identification of novel capabilities with biotechnological importance (Prista et al. 2016). A study conducted by Navarrete *et al*. (2020) demonstrated a beneficial role of salts, with sodium exerting a more significant effect, on the cell performance of *D. hansenii*, validating the role of salt in the yeast’s survival and establishing its halophilic behaviour. Simultaneously, their results suggested a positive and summative effect of acidic environment and high salt on the cell growth (Navarrete et al. 2020a).

The present study aims to establish an intraspecies behavioural characterisation of novel *D. hansenii* strains, through high-throughput screening methods using advanced robotics and automation devices. Micro-fermentation experiments are used to investigate the different strains’ capacity to tolerate high sodium concentrations and pH oscillations. Furthermore, carbon assimilation and growth inhibition induced by side-products from conventional feedstocks, is investigated using advanced screening robotics in solid media.

## Materials and Methods

### Strains and Culture Conditions

The strain library was obtained from the IBT culture collection of fungi from the Technical University of Denmark.

All strains were obtained from cryostocks (16% glycerol, Sigma-Aldrich, Germany) and were grown on yeast extract peptone dextrose (YPD) medium plates (10 g/L yeat extract, 20 g/L peptone, 20 g/L dextrose) containing 2% agar and at 28□.

Strains were cultivated in synthetic complete solid medium (6.7 g/L Yeast Nitrogen Base (YNB) w/o amino acids (Difco), 0.79 g/L complete supplement mixture (Formedium), 20 g/L dextrose) or YPD solid medium containing 2% agar. The pH was adjusted to 6.0 with NaOH. When needed, NaCl was added at the desired concentration (1M, 2M and 3M). All media were autoclaved at 121□ for 20 min. Separately filtered carbon source (glycerol, D-xylose, D-(+)-sucrose, D-arabinose, D-galactose, D-lactose, D-(+)-maltose, D-(+)-glucose) was added to the media to a final concentration of 2%. Inhibitors, furfural (1, 2, 4 g/L), hydroxy-methyl furfural (HMF) (0.5, 1, 2 g/L), vanillin (0.1, 0.2, 0.4 g/L), and methanol (1, 2, 4, 6 % v/v) were separately sterilized as well and added to the media when desired. Unless otherwise stated, cultures were incubated at 28□.

### Robustness Assay

The strain library was spotted from liquid to solid YNB medium, containing 2% carbon source and with/wo salt, using the ROTOR HDA (Singer Instruments, UK). When YPD solid medium was used, growth inhibitors in the desired concentrations were added, also in the presence or absence of salt.

### Microscopy Analysis

Qualitative cell growth was monitored after 48 h and 144 h using an in-house build digital camera setup (Nikon D90 SLR camera) or the PhenoBooth instrument (Singer Instruments, UK).

### High-throughput Micro Fermentations

Strains were pre-cultured in flasks containing YNB medium (pH 6), at 28□ and 250 r.p.m. for at least 24 h. Then, the cells were washed with ddH_2_O and re-inoculated in a 48-well FlowerPlate^®^ (MTP-48-B, m2p Labs, Germany) in YNB at pH 4 or 6, and with or w/o 1 M NaCl. The working volume in the plate was 1.5 mL at an initial OD_600_ of 0.1. The fermentations were performed in the BioLector® I (m2p Labs, Germany), adjusted to the following settings: temperature; 28□, filter; biomass (gain 10), humidity; on (>85% via ddH_2_O), oxygen supply; 20.85% (atmospheric air) and agitation speed; 1000 r.p.m. Light scattering units (excitation wavelength of 620 nm) were continuously monitored for at least 70 h. Data obtained from the micro-fermentation experiments were analysed using R version 4.0.3 (The R Foundation for Statistical Computing).

### Statistical Analysis

#### Heat Maps of Spot Tests

Strain’s growth on plates with or without 1M NaCl, was examined using the PhenoBooth instrument (Singer Instruments, UK) and belonging software. Following a uniform distribution, heat maps were generated based on standard scores (z-scores). Z-scores were obtained by (i) means of the control colony sizes, calculated on those strains cultivated in YPD plates containing inhibitors in the desired concentration (duplicates; n = 2), (ii) for each strain, measuring the distance to the mean on strains cultivated on the corresponding plates supplemented with 1M NaCl. Z-scores were illustrated in heat-maps with a purple to yellow spectra, corresponding to negative to positive values.

#### Analysis of Growth Rate

Background-subtracted light scattering units (LSU) were calculated by average measurements of uninoculated wells. Specific growth rates were calculated from the background-subtracted LSU and represented as the mean of the replicates, including a standard deviation. Variations in maximum specific growth rates (μmax (h^−1^)) were assessed with an analysis of variance (ANOVA test) using R (R Core Team 2020; Champely 2020). Local growth rates were estimated for each background-subtracted LSU by fitting a linear regression in rolling windows of six neighbouring measurement points, equivalent to estimating growth rates over a span of 18 minutes (Rugbjerg et al. 2018; Wickham et al. 2020; Zeileis and Grothendieck 2005). A moving regression based on a locally estimated scatterplot smoothing was applied to obtain scatterplot smoothing and to obtain a pattern in the progression of local growth rate. A confidence interval of 0.95 was included. The computation of local growth rates and the moving regression were constructed using (R Core Team 2020).

## Results

### Carbon Utilisation and Robustness Indications of *D. hansenii*

To assess robustness and investigate the carbon source utilisation and the effect of sodium in the different *D. hansenii* strains, we conducted a high-throughput spot-screening on selected carbon sources (Figure 1). While all strains (n=61) were able to utilise the majority of carbon sources investigated, weakened and varying growth was observed on both arabinose and xylose (on control plates and when supplemented with 1M NaCl) (Figure 1). In accordance with previous results obtained by Nobre *et al*. (1999), co-presence of medium-high salinity (1M sodium) and xylose or arabinose, indicates a tendency of improved carbon utilisation, and hence improved proliferation (Nobre, Lucas, and Leão 1999). Respectively, 1.7- and 8.2 percent points (pp) increase in strains able to grow on xylose or arabinose, were observed after 144 h of cultivation between control plates (no sodium) and plates supplemented with 1M sodium (data not shown).

**Figure 1.**
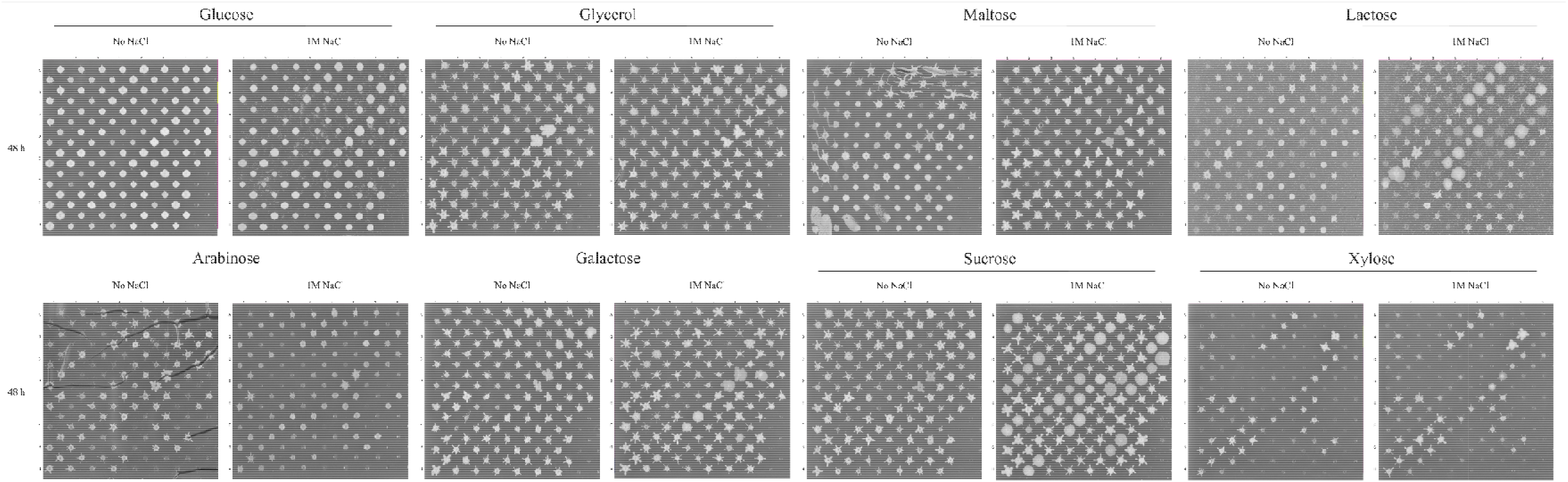
High-throughput spot-screening of carbon assimilation and effect of sodium on *Debaryomyces hansenii*. Cells were grown in YNB plates with or without 1M NaCl, and captures were obtained after 48h of cultivation. Position A1 contains *D. hansenii* CBS767 (a full map of all positions and strains can be found in Supplementary Table 1).

The percentage of carbon source assimilation by *D. hansenii* was estimated as the presence of growth after 48 h, and calculated from the spot-screening in YNB plates (Table 1). In the absence of salt, *D. hansenii* was successfully able to use all the sugars tested, being arabinose and xylose the ones with the lower estimated assimilation (78.7% and 67.2% respectively). The addition of 1M NaCl to the medium, seemed to improve the assimilation of arabinose but not xylose.

**Table 1.**
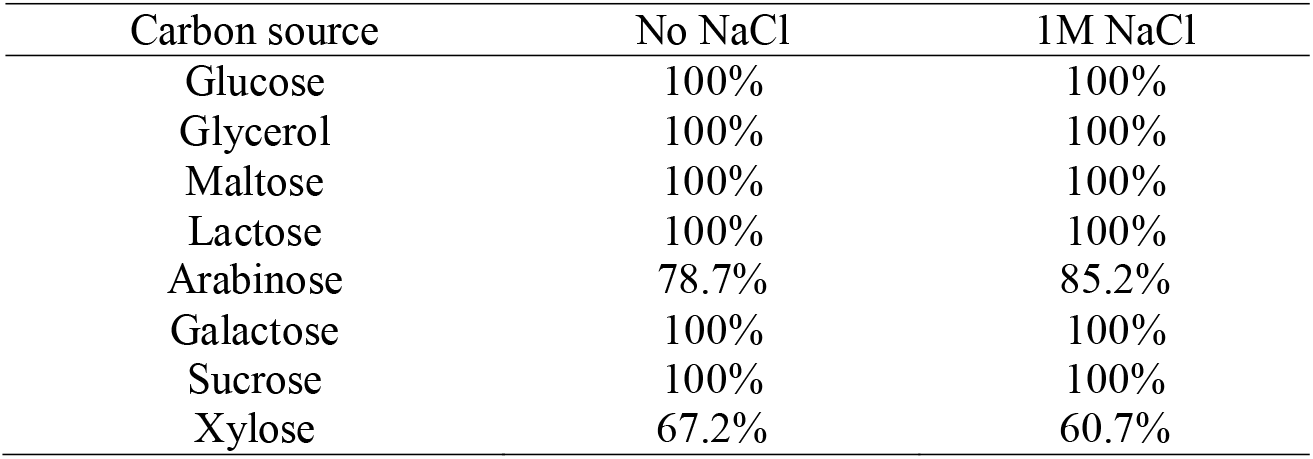
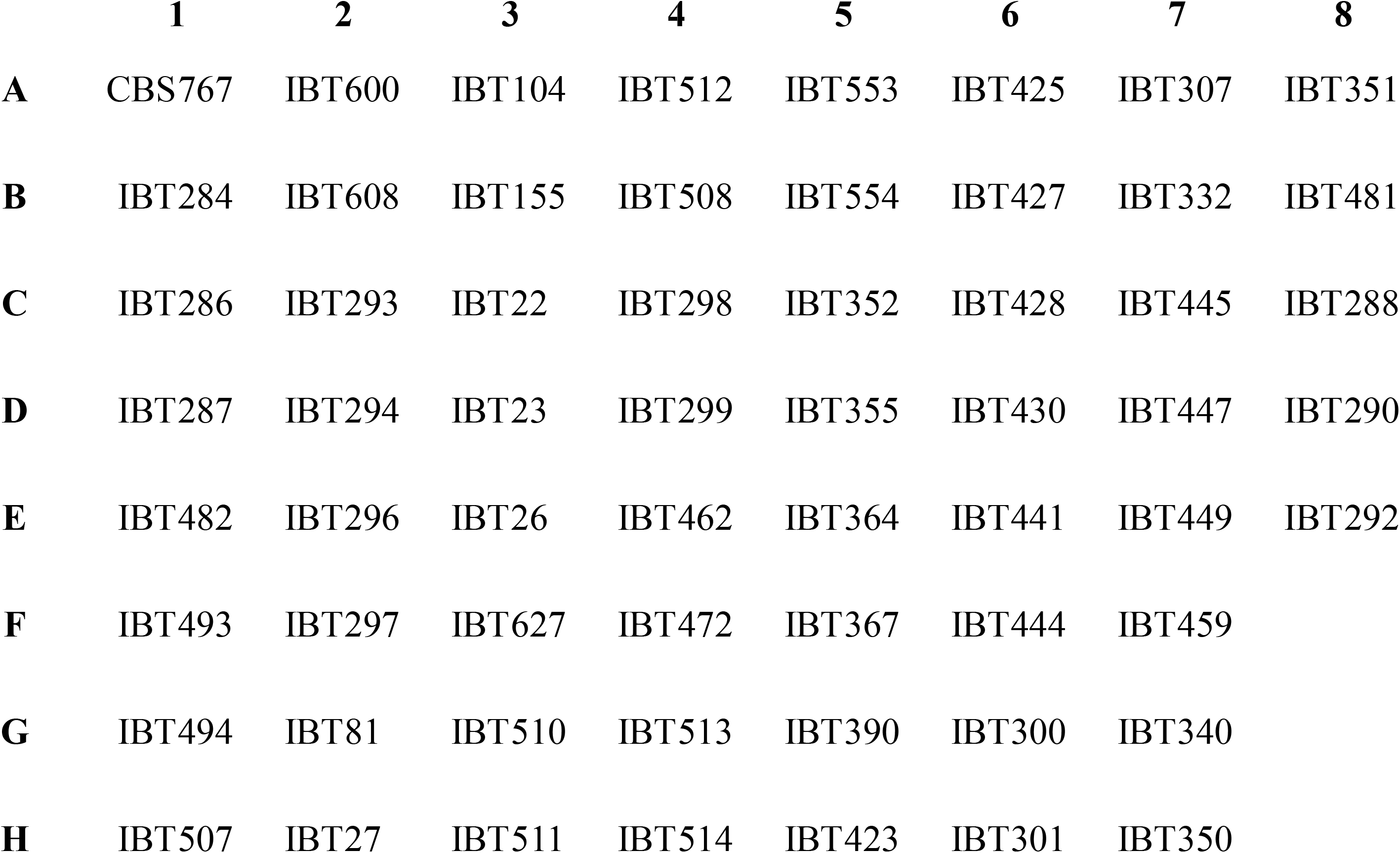
Carbon source utilization and effect of sodium on *Debaryomyces hansenii*, calculated from the spot-screening in YNB plates. Percentage of assimilation of carbon source was estimated as presence of growth after 48 h (n=2).

These results support the general assumptions and indications of non-inhibitory and non-detrimental effects of sodium on *D. hansenii* (Prista et al. 2016), while demonstrating that the yeast’s behaviour is not strain dependent, but it is extended intraspecies.

### Proliferation Capacity of *D. hansenii* at Higher Salinity

The general non-detrimental effect of 1M sodium observed in the spot-screenings for carbon utilisation (Figure 1 and Table 1), was further confirmed when we investigated the effect of higher concentrations of sodium (2M and 3M) on *D. hansenii’*s performance. Whilst 3M of NaCl detrimentally affected all isolates of *D. hansenii*, only slower growth was observed in the presence of 2M, as previously reported by Navarrete *et al*. (2020) (data not shown).

### Effect of Sodium on Cell Proliferation Capacity in the Presence of Inhibitory Compounds

To address the potential use of waste-derived feedstocks in bioprocesses with *D. hansenii*, we analysed co-presence of abundant inhibitory by-products from pre-treatment of both lignocellulosic and non-lignocellulosic materials (Kim 2018), and in the presence of sodium (Figure 2). Indications of a protective effect of sodium on the proliferation of *D. hansenii*, was observed when 2 g/L of furfural or 1 g/L of HMF were added to the medium. After 48 hours of cultivation, the colonies exhibited better growth on plates with both the inhibitor and salt, when compared to plates only containing the inhibitor. Similar results were obtained when half and double concentrations of furfural and HMF were used, and after 48 hours of cultivation (z-scores > 0) (Figure 2).

**Figure 2.**
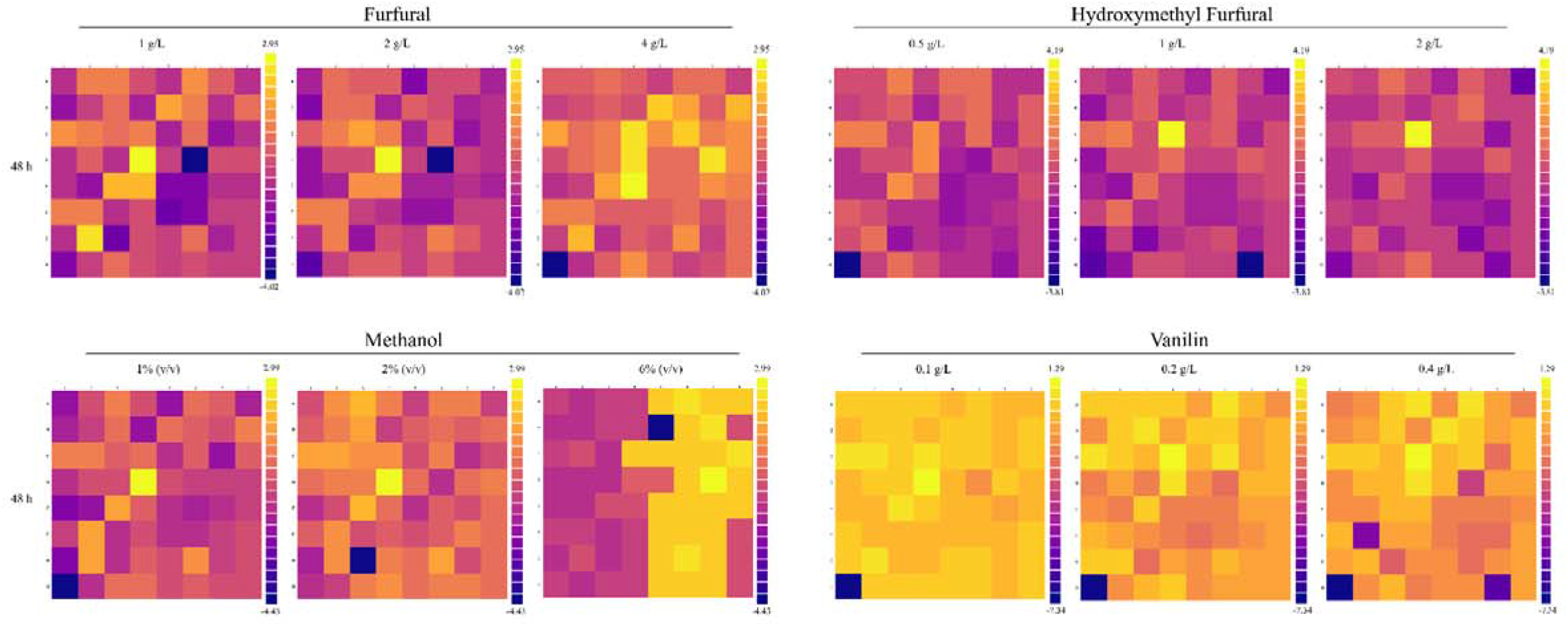
Heat-maps portraying z-scores of *Debaryomyces hansenii* strains (n=61) on 1M NaCl YNB plates (pH 6) including; furfural (1-, 2-, 4 g/L), hydroxymethyl furfural (HMF) (0.5-, 1-, 2 g/L), methanol (1-, 2-, 4\% (v/v)), and vanillin (0.1-, 0.2-, 0.4 g/L). Colony size of the corresponding *D. hansenii* strains on YNB plates (pH 6) including the aforementioned inhibitors constitute the plate mean. Heat-maps were obtained after 48h of cultivation using the PhenoBooth instrument and software. Differences in colony size are illustrated as a result of estimated z-score between the control plates (no sodium added) and plates containing inhibitors. Position A1 corresponds to *D. hansenii* CBS767 (a full map of all positions and strains can be found in Supplementary Table 1. A purple to yellow spectra corresponds to negative to positive z-scores and hence, worse to better capability to grow in the studied conditions.

While analysis of co-presence of sodium and the inhibitors methanol or vanillin also portrayed a non-detrimental and non-inhibitory effect of sodium, the supplement of 1M sodium did not appear to have a convincing engaging effect on cell proliferation after 48 hours of cultivation (Figure 2). This lack of effect was also observed at a lower concentration of methanol (1% (v/v)), where z-scores were distributed around zero and thus, suggesting little to no difference in the diameter span. 4% (v/v) of methanol seemed to affect cell proliferation of *D. hansenii* similarly to the observed effect at 2% (v/v), regardless the presence of 1M sodium in the medium (data not shown). At 6% (v/v) of methanol, a shift towards positive z-scores was observed after 48 h of cultivation. Thus, implying the presence of 1M of NaCl to have an enhancing effect on the cell proliferation capacity of *D. hansenii* at higher concentrations of methanol (Figure 2). On the other hand, the simultaneous presence of sodium and 0.1-0.2 g/L of vanillin, appeared to hamper cell proliferation. When colony sizes of *D. hansenii* cultivated on plates containing vanillin were compared to those on plates supplemented with 1M NaCl, z-scores were found to be negative. A varying tendency in cell growth was observed at 0.4 g/L of vanillin (Figure 2).

### Micro-fermentation Screening Analysis to Assess Effect of Sodium under Different Abiotic Parameters

A selection of the initial *D. hansenii* library was then analysed in the micro-fermentation screening analysis. Strains were selected based on (1) ability to utilise a broad spectrum of carbon sources, (2) substratum from which they were isolated, (3) capacity to grow in the presence of inhibitors

To address intraspecies behavioural trends of *D. hansenii* in 1M sodium and under different pHs (4 and 6), we studied biomass formation in simultaneous micro-fermentations experiments. The applied setup allowed for a semi-controlled environment and real-time monitoring of the growth of the selected strains. Scattered light signal, which is used as an indicator for biomass formation, may actually cause growth deviations, due to i.e. morphological changes associated with heterogeneity in the cell division cycle. Additionally, high cell densities frequently cause incorrect high measurements of biomass through the course of the bioprocess. These deviations are non-correctable (Kunze et al. 2014). Therefore, only maximum specific growth rates restricted to the exponential growth phase were used to compare growth between strains (Table 2).

**Table 2.** Maximum specific growth rates (μmax (h-1)) from micro-fermentations of selected *Debaryomyces hansenii* strains. Values were determined by light scattering units (LSU) obtained from batch cultivations in complete synthetic medium containing 2% glucose at 28□ and with initial pH of 4 or 6 (without pH regulation). (n=2).

| Strain | Condition | pH6<br>$\mu_{\max}$ (h<sup>-1</sup>) | pH6 (1M NaCl)<br>$\mu_{\max}$ (h<sup>-1</sup>) | pH4<br>$\mu_{\max}$ (h<sup>-1</sup>) | pH4 (1M NaCl)<br>$\mu_{\max}$ (h<sup>-1</sup>) |
| --- | --- | --- | --- | --- | --- |
| CBS767 |  | 0.226 | 0.231 | 0.060±0.009 | 0.102 |
| IBT22 |  | 0.126±0.029 | 0.130±0.021 | 0.049±0.003 | 0.050±0.006 |
| IBT23 |  | 0.210±0.020 | 0.187±0.031 | 0.081±0.001 | 0.086±0.018 |
| IBT26 |  | 0.146±0.019 | 0.227±0.018 | 0.064±0.025 | 0.069±0.017 |
| IBT27 |  | 0.308±0.036 | 0.412±0.029 | 0.131±0.010 | 0.155±0.014 |
| IBT81 |  | 0.097± | 0.243±0.024 | 0.042 | 0.054±0.010 |
| IBT155 |  | 0.299±0.006 | 0.146±0.073 | 0.017 | 0.027±0.001 |
| IBT290 |  | 0.238±0.001 | 0.195±0.010 | 0.015±0.002 | 0.082±0.003 |
| IBT350 |  | 0.251±0.020 | 0.145±0.043 | 0.032±0.001 | 0.059 |
| IBT364 |  | 0.353±0.010 | 0.278±0.038 | 0.013±0.002 | 0.072±0.004 |
| IBT367 |  | 0.271±0.010 | 0.385±0.023 | 0.075±0.012 | 0.091±0.010 |
| IBT427 |  | 0.203±0.026 | 0.180±0.008 | 0.099±0.004 | 0.141 |
| IBT444 |  | 0.280±0.034 | 0.302 | 0.071±0.001 | 0.074±0.004 |
| IBT447 |  | 0.151 | 0.249±0.004 | 0.051±0.004 | 0.061 |
| IBT449 |  | 0.234±0.062 | 0.235±0.008 | 0.070±0.007 | 0.084±0.001 |
| IBT493 |  | 0.238±0.002 | 0.184±0.002 | 0.085±0.002 | 0.085±0.009 |
| IBT510 |  | 0.135±0.022 | 0.059±0.019 | 0.049 | 0.070 |
| IBT600 |  | 0.139±0.034 | 0.213±0.007 | 0.048±0.011 | 0.060 |
| IBT608 |  | 0.107±0.025 | 0.087±0.010 | 0.054±0.011 | 0.062 |
| IBT627 |  | 0.299±0.004 | 0.231±0.041 | 0.077±0.007 | 0.088±0.021 |

Differences in growth were observed in both the presence and absence of 1M sodium in the media. At pH 6, addition of sodium did not throw any conclusive results, as some strains showed an increased in growth while others did not.

Previous studies have shown synergistic and protective effect of NaCl when other stress factors are present. Specifically, a positive correlation between available Na^+^ and acidic environments is proved to enhance cell growth, in comparison to non-saline, acidic environments (Almagro et al. 2000; Navarrete et al. 2020a). On the other hand, and although a general differential growth was observed intraspecies at pH 4, a more conclusive positive effect on biomass formation was observed when 1M sodium was present (Table 2). Additionally, IBT27 strain reached the highest maximum specific growth rate in all the conditions studied. Importantly, these results, in accordance with previous published work, indicate a synergistic and protective effect of low pH and high salt on the cell growth of *D. hansenii* (Almagro et al. 2000; Navarrete et al. 2020a; Prista et al. 2016), and it is not strain-dependent.

### Local Growth Rates Measured During the Course of the Simultaneous Micro-fermentations

In order to study the dynamics of the strains’ fitness over time and to understand the influence of the abiotic parameters on their stability, we measured local growth rates throughout the micro-fermentation experiments. Generally, a stable biomass formation was observed in all the strains, when 1M NaCl was present in the media (Figure 3). Under non-acidic cultivation (pH 6) and without sodium present, some strains were able to reach higher maximum specific growth rates (e.g. strain IBT155, IBT364, and IBT493). However, a more pronounced decrease in local growth rate was observed over time in these specific cases, resulting in a lower average local growth rates. When cultivated in the presence of sodium, strains with higher average maximum specific growth rates, did now show an abrupt decrease in the local growth rate (e.g. strain CBS767, IBT22, and IBT81) (Figure 3). Growth-coupled production by selecting for cells with higher fitness is a desirable target for strain design (Dinh et al. 2018). In conclusion, higher fitness of the strains was not observed to be correlated with growth stability in the absence of sodium and at pH 6.

**Figure 3.**
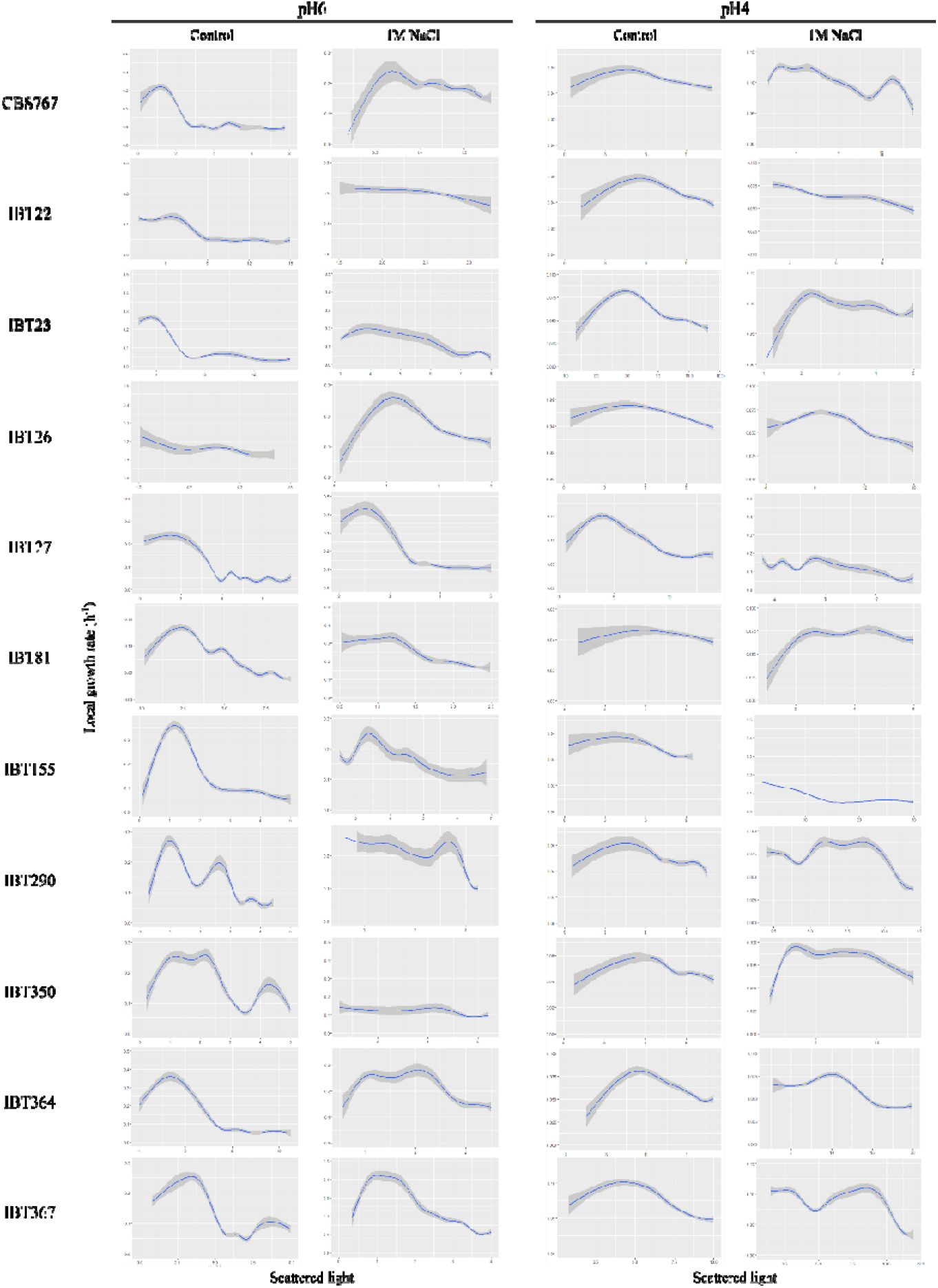
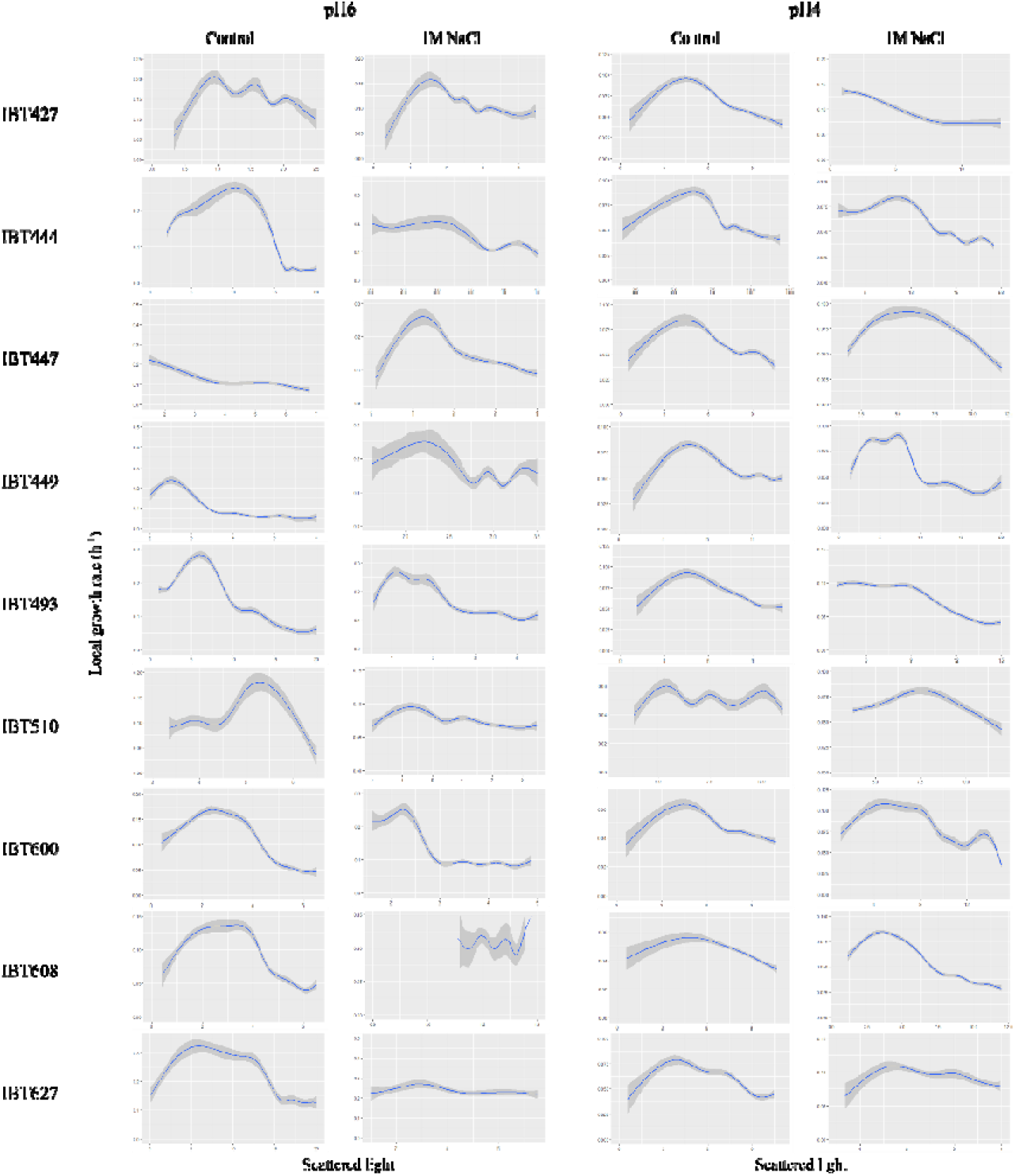
Local growth rates for *Debaryomyces hansenii* strains in YNB medium (pH 6 and pH 4) supplemented with 1M NaCl. Local growth rates were measured throughout the course of the micro-fermentation experiments. Smoothed conditional means were calculated using locally estimated scatterplot smoothing. Light grey areas represent confidence interval of 0.95 (n = 2).

When the strains were cultivated at acidic pH (pH 4) and in the presence of sodium, a more stable progression in growth throughout the course of the experiments was observed (Figure 3). Interestingly, these conditions also promoted higher maximum specific growth rates for all the strains, when compared to pH 4 without sodium (Table 2). Once more, the observed improvement in growth indicates that the synergistic and protective effect of low pH and high sodium on cell growth, is not strain dependent, but a general tendency within the *D. hansenii* species (Almagro et al. 2000; Navarrete et al. 2020a; Prista et al. 2016).

## Discussion

The global societal transition from chemical, conventional production of bulk- and high value molecules to sustainable solutions is dependent on innovative bio-based production. Developing cost-efficient solutions, will help to obtain the maximal potential of industrial biotechnology, and thus, to accelerate the transition to a more sustainable processes (Erickson, Nelson, and Winters 2012). The non-conventional yeast *D. hansenii* presents some important physiological properties that have generated an increased interest in this yeast as a promising host in bioprocesses (Prista et al. 2016). Its unique and prominent halotolerant characteristics ensure its importance in industrial applications (Chao, Yen, and Ku 2009; Prista et al. 2016).

While the halotolerant behaviour of *D. hansenii* was already established for several authors (Lépingle et al. 2000; Souciet et al. 2009), recent studies have revealed the salt-including and halophilic behaviour of this yeast (Navarrete et al. 2020a; Prista et al. 2016). Similar conclusions are obtained from the present study. Medium-high concentration of sodium (1M) does not show a detrimental effect on the carbon assimilation capacity of any of the strains included in this study. From our results, *D. hansenii’s* ability to assimilate a broad spectrum of carbon sources (Nobre, Lucas, and Leão 1999) can be now extrapolated to other strains of the same species (Figure 1). A varying growth was observed in the presence of arabinose and xylose in the different strains tested, which is in alignment with previous work reporting slower and weaker growth of *D. hansenii* when assimilating pentoses (Nobre, Lucas, and Leão 1999; Palma et al. 2007; Pereira et al. 2014) Moreover, an increase in the capacity to assimilate pentoses was observed with the addition of 1M sodium, highlighting again the halophilic behaviour of *D. hansenii* strains.

A very recent study conducted by Navarrete *et al*. (2020), demonstrated an increased growth in the presence of up to 1M NaCl when compared to control conditions (no salt). Therefore, non-detrimental and non-inhibitory effects of sodium on the proliferation capacity of *D. hansenii* were described by the authors. Despite the different applied method in this study, our results from the high-throughput spot-screenings support previous observations by Navarrete *et al*. (2020). An improved growth was observed when *D. hansenii* strains were cultivated in the presence of 1M sodium (Figure 1).

### Increased Stability of *D. hansenii* Strains when Cultivated in Acidic Culture Conditions

Whereas mutation-prone cells secure adaptation by shifting environmental changes for the evolving population, each underlying mechanism presents a probable threat to the stability of the introduced synthetic system. Evolving sub-cultures of non-producers, is an increasing issue in long-term cultivation cell factory strains (Hewitt and Nienow 2007). Generally, an improved stability of the library is observed when sodium is present in the culture media, during the micro-fermentations (Figure 3). Importantly, the stability is even more profound when cells are cultivated in acidic culture conditions. In addition, an initial higher maximum specific growth rate was observed for all the strains in pH 4 and medium supplemented with 1M sodium (Table 2). A summative effect of acidic environments and high salt content has previously shown to have a positive influence on cell growth (Almagro et al. 2000; Navarrete et al. 2020). Our results further sustained previous observations as all the strains of *D. hansenii* tested in this study showed higher maximum specific growth rates in acidic environments supplemented with sodium (Table 2). Subsequently, more stable local growth rate profiles were also observed at low pH and in the presence of sodium (Figure 3).

Interestingly, strain IBT27, which showed broad carbon source assimilation and high resistance to inhibitors, especially in the presence of 1M NaCl, also reached the highest maximum specific growth rate at low pH and high sodium (Figure 1 and Table 1). Consequently, IBT27 can be considered as one of the best candidates among the other tested strains. However, at pH 6 and with 1M NaCl, the maximum specific growth rate was lower compared to that with no sodium added. Therefore, our results align very well with previous work showing an improved performance of *D. hansenii* in environments under the presence of abiotic stresses, and a positive effect on growth in acidic environments with high sodium content (Almagro et al. 2000; Navarrete et al. 2020a). Additionally, we present strong indications of strain-specific responses to the presence of sodium, with some strains responding much better to high sodium than the standard strain (CBS767).

The use of high throughput technology facilitates the screening of a range of different parameters in a reasonable short period of time, while significantly contributing to understand the complexity of bioprocesses in general. In addition, the type of analysis used in this work allows the detection of significant differential behaviour between strains at a colony-growth level. Hence, this study provides sufficient prove to be employed in future screenings and identification of optimal candidates of engineered *D. hansenii* cells, e.g. to express a product of interest.

### *D. hansenii’* Response to Stresses derived from Lignocellulosic Biomass and Non-lignocellulosic Feedstocks

Abundant and renewable lignocellulosic biomass has high potential as a substitute for fossil resources to produce bioprocessed chemicals and materials, while eluding less available and more expensive feedstocks (Li et al. 2017). However, pre-treatment of lignocellulosic biomass yields several inhibitory by-products, making the use of lignocellulosic biomass as feedstock problematic in some cases (Navarrete et al. 2020b). Inhibitory compounds from biomass pre-treatment processes include furan derivatives compounds (including furfural and HMF) and phenolic compounds, such as vanillin (Ask et al. 2013). We found sodium to relieve the abiotic stress caused by the presence of different concentrations of furfural and HMF (Figure 2). This reinforces previously results suggesting that sodium does not have a toxic effect on *D. hansenii* cells, and being able to protect the yeast cells against additional abiotic stress factors (Ramos-Moreno, Ramos, and Michán 2019; Navarrete et al. 2020a).

Furan derivatives were found to severely affect the redox metabolism of *S. cerevisiae* while lower inhibition of the energy metabolism was observed (Ask et al. 2013). A study conducted by Ramos-Moreno *et al*. (2019) demonstrated that expression of genes related to oxidative stress were induced in *D. hansenii* by high salt concentrations (with sodium enabling a better induction than potassium). Hence, suggesting a direct link between high salt concentrations and ROS detoxification. Moreover, a very recent study conducted by Navarrete *et al*. (2020) demonstrated an increased respirative activity of *D. hansenii* in the presence of salts, especially sodium. The ability of sodium to enhance the respirative metabolism and to induce expression of ROS detoxification genes (Ramos-Moreno, Ramos, and Michán 2019; Navarrete et al. 2020a) may explain why the proliferation capacity of *D. hansenii* strains is inhibited by furan derivatives, when no sodium is available, but improved when sodium is present.

On the contrary, the simultaneous presence of sodium and vanillin, known to inhibit the activity of the DNA-dependent protein kinase catalytic subunit, appears to have an opposite effect, resulting in hampered cell proliferation in *D. hansenii* strains (Figure 2). However, depending on the lignocellulosic biomass composition and pretreatment used, a vast range of vanillin concentrations can be present, from 1-26 mM (Ito et al. 2020). Therefore, the effect of higher concentration of vanillin should be further investigated in order to get a better picture about the inhibitory impact of vanillin on *D. hansenii* strains. In addition, the observed summative effect of sodium on cell proliferation in the presence of furfural and HMF should be also further investigated, including both furan derivatives and phenolic compounds.

An improved proliferation capacity was also observed at high concentrations of methanol (Figure 2). The results obtained from the spot-test suggest a robust behaviour of all *D. hansenii* strains, with some of themoutperforming the average.

Finally, our study sheds light on the behavioural patterns of novel *D. hansenii* strains in semi-controlled micro-fermentations and spot-test analysis, which contribute to establish the fundamental knowledge about this non-conventional yeast. A successful implementation of industrial scale bioprocesses is restricted by limited access to cost efficient solutions (Li et al. 2014). From the inhibitors study presented in this work, it was found that lignocellulosic biomass and non-lignocellulosic feedstocks could be used in *D. hansenii*-based industrial bioprocesses, which will decrease the process costs while increasing the potential use of this yeast in the market. Furthermore, culture media or water supply containing high salt concentration (including sea-water) as well as open continuous bioprocesses could be used in industrial set-ups, due to the halophilic character of this yeast, and so bypass the need for extensive sterilisation processes.

## Conclusion

The use of HT-technology to monitor the different behavioural patterns of several *D. hansenii* strains was successfully used in the present study. Besides, our findings align very well with previous published work of this yeast.

An improved performance in the presence of abiotic stresses was observed when 1M NaCl was added to the media. Significantly, a positive and summative effect on *D. hansenii* strains’ fitness was observed when pH 4 and high sodium was used. Moreover, our results support previous reports describing a protective effect of sodium on *D. hansenii’*s growth, further expanded to the use of inhibitors that are released from pre-treatment of lignocellulosic on non-lignocellulosic feedstocks. This reveals the great potential of *D. hansenii* in different industrial applications and bioprocesses.

Importantly, while some level of differentiation was observed within the different strains tested, the halophilic behaviour and sodium-induced optimal performance of *D. hansenii* can be now extended to all strains included in this study.

